# Alpha-synuclein overexpression can drive microbiome dysbiosis in mice

**DOI:** 10.1101/2024.02.01.578464

**Authors:** Timothy R. Sampson, Zachary D Wallen, Woong-Jai Won, David G. Standaert, Haydeh Payami, Ashley S. Harms

## Abstract

Growing evidence indicates that persons living with Parkinson disease (PD), have a unique composition of indigenous gut microbes. Given the long prodromal or pre-diagnosed period, longitudinal studies of the human and rodent gut microbiome prior to symptomatic onset and for the duration of the disease period are currently lacking. PD is characterized in part by accumulation of the protein α-synuclein (α-syn) into insoluble aggregates, in both the central and enteric nervous systems. As such, a number of experimental rodent and non-human primate models of α-syn overexpression recapitulate some of hallmark pathophysiologies of PD. These animal models provide an opportunity to assess how the gut microbiome changes with age under disease relevant conditions. Here, we used a transgenic mouse strain, the Thy1-hSYN “line 61” mice which over express wild-type human α-syn to test how the gut microbiome composition responds in this model of PD pathology during aging. Using shotgun metagenomics, we find significant, age and genotype dependent bacterial taxa that become altered over age. We reveal that α-syn overexpression can drive alterations to the gut microbiome composition and suggest that it limits the expansion of diversity through age. Given emerging data on potential contributions of the gut microbiome to PD pathologies, our data provide an experimental foundation to understand how the PD-associated microbiome may arise as a trigger or co-pathology to disease.

## INTRODUCTION

Parkinson disease (PD) is a common, progressive, neurodegenerative movement disorder. Pathologically, PD is characterized by loss of dopamine-producing neurons in the substantia nigra pars compacta (SNpc) and neuronal accumulation of pathological forms of the protein α-synuclein (α-syn) into insoluble aggregates, termed Lewy bodies. This accumulation of α-syn results in numerous cellular and immune dysfunctions and eventually neuronal cell death in the SNpc, as well as in other brain regions. While most cases of PD are idiopathic, genetic mutations or duplications in the genes encoding α-syn, *SNCA*, which increase expression or result in the disruption of the native structure of α-syn, lead to familial forms of PD (*1*). Genome-wide association studies (GWAS) show strong associations between variants in or near *SNCA* that affect expression of α-syn and increased risk of sporadic PD (*2, 3*). Various experimental rodent and non-human primate models of α-syn overexpression recapitulate some of these physiologic and pathologic characteristics (*4, 5*). Collectively, these data point to a central role for α-syn in the etiology of PD, but the mechanisms by which α-syn initiates or drives disease pathogenesis are unknown.

Gastrointestinal (GI) dysfunction is a prevalent prodromal symptom of PD observed either preceding and/or concurrent with diagnostic motor symptoms. Common symptoms such as gastroparesis and chronic constipation persist for the duration of the disease, which impacts patient quality of life. While α-syn aggregation in the basal ganglia and/or cortical regions of the central nervous system (CNS) is one pathological hallmark of PD, α-syn pathology and inclusions are also reported in the olfactory bulb, brainstem, and the enteric nervous system (ENS). Therefore, PD is not just a disease of the CNS, but may involve the ENS and GI tract prior to overt neurodegeneration and movement symptoms. The pattern of α-syn pathology led Braak et al. to hypothesize that abnormal α-syn pathology may begin in the gut and propagate via a prion-like fashion into the CNS via the vagus nerve(*6*). This “gut to brain” hypothesis is supported by α-syn+ inclusion bodies in the ENS and vagus nerve in post-mortem tissues (*6, 7*). Further, the risk of developing PD was significantly reduced among individuals that received a full truncal vagotomy(*8*). Experimental support of this hypothesis is observed in preclinical rodent models. Injections of α-syn pre-formed fibrils directly into the GI tract leads to pathology in the CNS in rodents(*9-11*), also observed in similar experiments involving non-human primates (*12*).

The majority of PD incidences are idiopathic and likely involve genetic and/or environmental interactions. It is possible that factors intrinsic to the GI tract, such as the gut microbiome, may contribute to the etiology or progression of PD pathogenesis. The gut microbiome consists of trillions of microbes that inhabit the GI tract and play important physiological roles across many aspects of health and disease. Numerous studies to date have highlighted compositional differences in the gut microbiome between persons with PD compared to neurologically healthy controls including detection of altered abundances in several microbial species (*13-15*). Experimental models have demonstrated varying levels of contribution by the gut microbiome to aspects of PD pathology including both α-syn dependent and toxicant-induced model systems(*16-20*). Whether the gut microbiome is directly involved in the etiopathogenesis of PD in humans is unknown.

While experimental studies suggest the composition of the gut microbiome can impact α-syn-mediated disease pathogenesis, it remains unclear how PD-associated dysbiosis of the gut microbiome arises in an individual. Elucidating how the gut microbiome is first impacted in those with PD and subsequently shifts over disease progression is key for understanding how these experimental contributions ultimately relate to disease. One possibility is that underlying α-syn-mediated dysfunctions lead to an altered intestinal environment which impacts the gut microbiome composition. In the current study, we sought to test three major outcomes: (1) determine whether α-syn overexpression impacts gut microbiome composition in mice, (2) establish whether this dysbiosis is present from birth or develops with age, and (3) identify the specific microorganisms that likely drive this dysbiosis. We longitudinally sampled fecal pellets for 12 months from transgenic (TG) mice overexpressing α-syn under a Thy1 promoter (Thy1-SYN “line 61” TG mice) (*21*) and their non-transgenic (NTG) littermate controls and performed shotgun metagenomics followed by taxonomic profiling. Our analysis revealed significant, age dependent changes in the gut microbiome of TG mice compared to NTG control mice, indicating α-syn overexpression alone can drive dysbiosis *in vivo*. Differential abundance analysis of individual species revealed a highly significant 10 to 50-fold reduction in the abundances of *Lactobacillus* and *Bifidobacterium* species, taxa that are conversely and consistently found highly enriched in PD (*13, 15*). The results from this study indicate α-syn overexpression in mice drives significant, age dependent gut dysbiosis that correlates with ages at which hallmark model dependent motor and histological pathologies occur. These results suggest that while α-syn overexpression is sufficient to drive gut dysbiosis in mice, including taxa impacted in PD, it does not recapitulate the human PD-associated microbiome. Continued investigation into the parameters that lead to PD-associated gut dysbiosis are necessary to how this arises in PD as an etiological trigger or a co-pathology that contributes to immune and other PD-related phenotypes.

## METHODS

### Mice

Female mice overexpressing human α-syn under the Thy1 promoter (Thy1-SYN “line 61” TG female mice) originally developed in the laboratory of Elizer Masliah at the University of California San Diego (*21*), were bred to BDF1 males purchased from Charles River Laboratories (Wilmington, MA, USA) as previously described (*22*). As the Thy1-SYN transgene insertion site is on the X chromosome, male α-syn transgenic mice exhibit age-dependent α-syn pathology, neuroinflammation including gliosis, and motor symptoms. Despite PD-like phenotypes, α-syn TG mice do not develop dopaminergic cell loss or α-syn positive inclusions in the SNpc. Male and female mice were separated prior to genotyping 21 days post birth, and TG and NTG genotyped littermates were co-housed for the duration of the study. All research conducted on animals was approved by the Institutional Animal Care and Use Committee at the University of Alabama at Birmingham.

### Fecal pellet collection and processing

#### Analytical sample

Fecal pellets were collected at four time points (1, 3, 6, and 12 months) from 22 TG and 32 NTG mice across 22 litters (total analytical sample N = 216). Nine out of the 22 litters included both TG and NTG mice and were included in sensitivity analyses (N = 11 TG and 22 NTG littermates). Fecal pellets were counted and collected from individual mice over a 20 minute period in a novel clean cage environment by using sterile toothpicks, then immediately labeled and stored at -80°C until DNA extraction.

#### DNA isolation, library preparation and next generation sequencing

Isolation of fecal DNA, sequence library preparation, and next generation sequencing was performed at CosmosID Inc. (Germantown, MD, USA). Isolation of DNA from fecal pellets was performed using QIAGEN DNeasy PowerSoil Pro Kit (Germantown, MD, USA) according to the manufacturer’s protocol, then quantified using Qubit™ 4 fluorometer and dsDNA HS Assay Kit (Thermofisher Scientific, Waltham, MA, USA). Isolated DNA was stored at -20°C until library preparation. Sequence libraries were prepared using the Nextera XT DNA Library Preparation Kit (Illumina, San Diego, CA, USA) and Integrated DNA Technologies unique dual indexes (San Diego, CA, USA) with a total DNA input of 1ng. Genomic DNA was fragmented using a proportional amount of Illumina Nextera XT fragmentation enzyme. Unique dual indexes were added to each sample followed by 12 cycles of PCR to construct libraries. Sequence libraries were purified using AMPure magnetic beads (Beckman Coulter, Brea, CA, USA) and eluted in QIAGEN EB buffer. Sequence libraries were quantified using the Qubit™ 4 fluorometer and dsDNA HS Assay Kit. Libraries were then sequenced on an Illumina HiSeq X platform with 150 bp paired-end sequencing. Number of reads ranged from 1M to 5M per sample.

#### Bioinformatic processing of sequences

We used FastQC (https://www.bioinformatics.babraham.ac.uk/projects/fastqc/) and MultiQC (*23*) to check the quality of raw sequences, followed by processing of sequences using BBDuk v 38.92 (https://jgi.doe.gov/data-and-tools/software-tools/bbtools/) to remove adapters and PhiX sequences and quality trim and filter low-quality sequences. BBDuk was performed specifying ‘ftm=5’, ‘tbo’, ‘tpe’, ‘qtrim=rl’, ‘trimq=25’ and ‘minlen=50’ as described previously (*13*). Next, all sequence reads mapping to the most recent version of the Mus musculus genome reference (GCA_000001635.9_GRCm39_genomic.fna) were removed using BBSplit v 38.92 with default parameters. Low-complexity sequences were filtered using BBDuk specifying ‘entropy=0.01’. No sample was excluded due to low sequencing depth. Number of reads per sample ranged from 0.3M to 3.1M after processing.

#### Taxonomic profiling

Taxonomic profiling was performed for processed sequences using MetaPhlAn v 3.0.14 with default parameters (*24*) and resulted in the detection of 48 species from 29 genera, ranging from 5 to 22 species per mouse. Relative abundances of taxa (used for differential abundance testing, i.e. MWAS) were calculated using MetaPhlAn default settings. To calculate count data (used for beta- and alpha-diversity-based analyses), MetaPhlAn was ran again adding the ‘--unknown-estimation’ flag to generate relative abundances with unknown estimation (i.e., relative abundances that take the proportion of “unknown” microbes in a sample into account), then relative abundances were multiplied by the total sequence reads of the sample.

### Statistical analyses

For all statistical analyses and visualizations, R v 4.3.1 (https://www.r-project.org/) was used.

#### Beta- and alpha-diversity

To measure differences in gut microbiome composition between samples (beta-diversity), we used Aitchison distances. Aitchison distances were calculated for each time point by taking the Euclidean distances of centered log-ratio (clr) transformed species counts. The clr transformation was performed using the following formula in R:

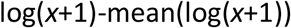

where *x* is a vector of all species counts in a sample. Aitchison distances were used as input to principal component analysis (PCA) and permutational analysis of variance (PERMANOVA)(*25*). PCA was performed to observe the beta-diversity among samples by first generating principal components (PCs) using the ‘prcomp’ function in R, then plotting PC 1 and 2 for each time point using the ‘autoplot’ function from the ggfortify v 0.4.16 R package (https://cran.r-project.org/web/packages/ggfortify/index.html). PERMANOVA was performed to test for differences in beta-diversity between TG and NTG groups for each time point using the ‘adonis2’ function from the vegan v 2.6.4 R package (https://cran.r-project.org/web/packages/vegan/index.html) specifying permuted P-values to be calculated with 9,999 permutations.

To measure intra-sample diversity of the gut microbiome (alpha-diversity), three diversity metrics were calculated (observed richness, Simpson index, Shannon-Weiner index) using the ‘diversity’ function in vegan. Differences in alpha-diversity between TG and NTG groups at each time point were tested using linear regression (via the ‘lm’ function in R) adjusted for the total sequence count per sample, and additionally, PC1 from PCA for 1, 3, and 6 months. Both total sequence count and PC1 were standardized using the ‘scale’ function in R prior to testing.

#### Differential abundance testing

To test for differentially abundant species and genera between TG and NTG groups, linear regression was performed on log2-transformed relative abundances as implemented in MaAsLin2 v 1.14.1 (*26*) for 32 species and 19 genera captured during the 12^th^ month of sampling. MaAsLin2 was ran using default parameters with the following exceptions: ‘min_prevalence’ was set to ‘0.05’ making the effective sample size N = 3, ‘normalization’ was set to ‘NONE’ as we are already inputting relative abundances (which are normalized by the total sequence count per sample), and ‘standardize’ was set to ‘FALSE’ as standardization was performed for quantitative variables prior to running MaAsLin2. P-values from differential abundance testing were multiple testing corrected separately for species- and genus-level comparisons using the Benjamini-Hochberg false discovery rate (FDR) method [PMID: 2218183]. Additionally, testing was performed for earlier timepoints (1, 3, and 6 months) for species that were found differentially abundant between 12 month TG and NTG in order to observe their differential abundance as a function of age. All differential abundance analyses were adjusted for total sequence count per sample, and analyses for 1, 3, and 6 month time points were additionally adjusted for PC1.

#### Sensitivity analysis

To ensure results were robust to any litter effects, as some litters contained only TG or NTG mice, all aforementioned analyses (beta- and alpha-diversity-based and differential abundance testing) were performed again including only litters that resulted in both TG and NTG litter mates (9 mouse litters out of 22). No difference in methodology was implemented save the fact that TG and NTG sample sizes were reduced by 50% (N = 22 to N = 11) and 38% (N = 32 to N = 20), respectively.

## RESULTS

Human observations strongly demonstrate that persons with PD harbor a gut microbiome that is compositionally different from healthy controls. Experimental data suggest contributions by certain gut bacteria to PD-relevant pathologies in rodent models. However, what drives these alterations to the intestinal environment and selects for these compositions is unknown. We therefore sought to determine whether α-syn overexpression, a pathological attribute associated with PD, is sufficient to induce gut dysbiosis in mice. We collected fecal pellets from 54 male mice, including 22 Thy1-SYN “line 61” mice (TG) and 32 non-transgenic littermate controls (NTG), at four consecutive time points: 1 month, 3 months, 6 months, and 12 months post-birth. Littermates provide for a near identical maternally derived microbiome between genotypes. DNA was extracted and shotgun sequencing was performed, generating an average of 1M to 5M sequence reads per sample before sequence QC and 0.3 to 3.1M after QC. All samples passed QC. We used standard protocols for bioinformatics (bioBakery suite), taxonomic profiling (MetaPhlan3), and statistics (MaAsLin3) to assess bacterial compositions.

### Visual inspection of microbiome similarities as mice age

We first extracted principal components (PC) from microbial abundance data and created PC distance plots for qualitative assessment of similarity/dissimilarity of microbiomes at each age independently (**Figure 1**). In mice aged 1 month, we detected an underlying structure (an unknown source of variation) which separated the samples into two clusters of 15 mice and 40 mice each. The first principal component (PC1) explained 42% of the total variation seen at month 1. This structure became weaker as the mice aged; with this cluster diminishing as mice aged. The visible clustering was dissolved completely by month 12. To control for this variation in early life, in downstream analysis, we adjusted for PC1 when testing ages 1, 3 or 6 months, but not 12 months. We do, however, note the appearance of a genotype effect that begins to define TG vs NTG-derived microbiomes by 12 months of age (**Figure 1**).

**Figure 1.**
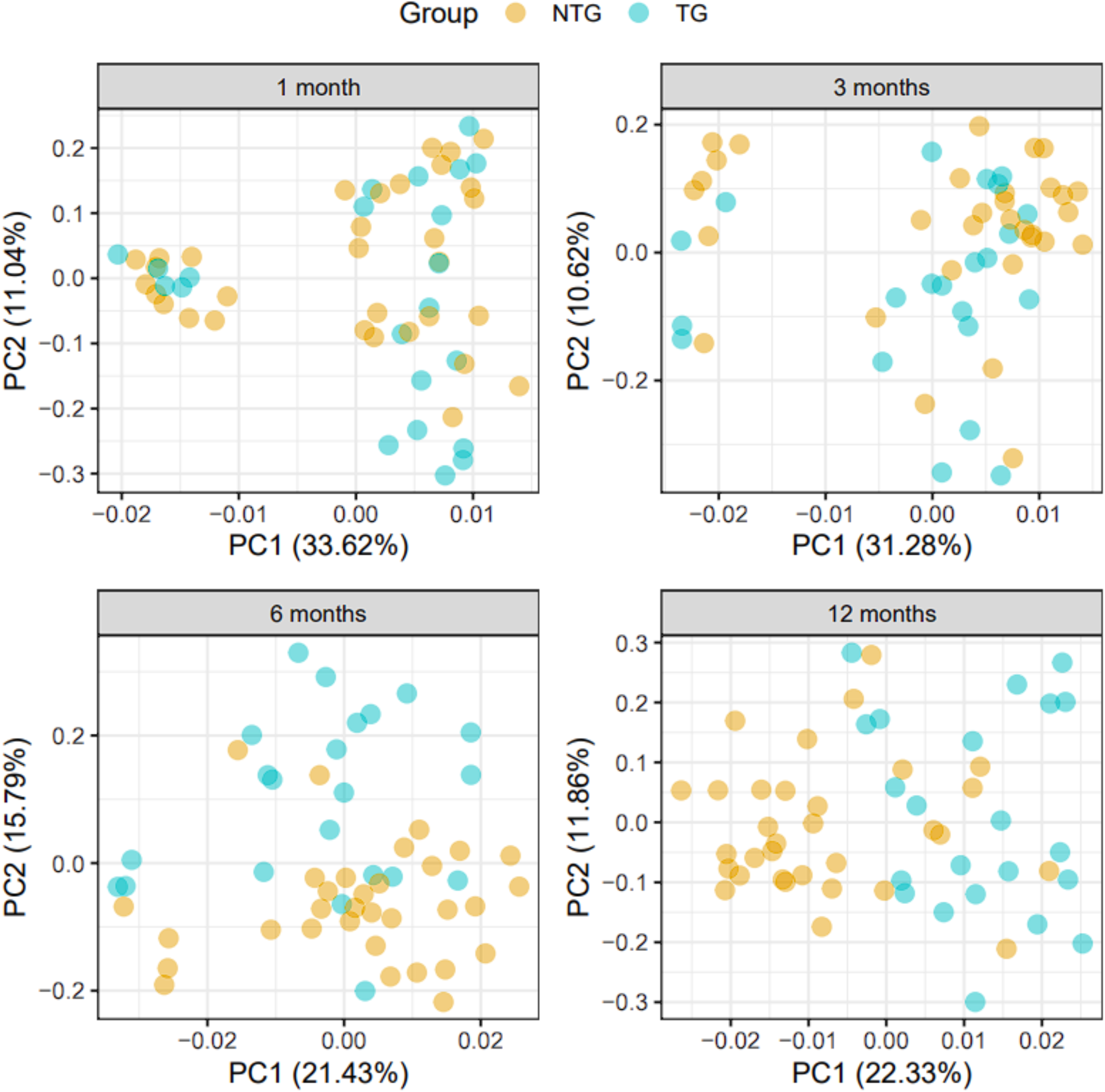
Inter-sample differences in gut microbiome composition (beta-diversity) of Thy1-SNCA transgenic and non-transgenic mice at 1, 3, 6 and 12 months. Principal component (PC) analysis was performed to visualize inter-sample differences in gut microbiome composition (beta-diversity) between Thy1-SNCA transgenic (TG, blue points, N = 22) and non-transgenic (NTG, orange points, N = 32) mice at each time point. Each point in the plots represents the composition of the gut microbiome of one unique mouse sample at a certain time point and distances between points indicate degree of similarity of a mouse sample to others. Percentages on the x- and y-axis correspond to the percent variation in gut microbiome compositions explained by PC1 and PC2. The differences between TG and NTG groups for each time point were formally tested using PERMANOVA (Table 1a).

### α-syn overexpression induces dysbiosis in the gut

We therefore conducted a formal statistical test to determine if the overall composition of gut microbiome in TG mice was different from NTG mice **(Table 1a)**. We tested the Atchison distances between samples (beta-diversity) using permutational multivariate analysis of variance analysis (PERMANOVA) to find whether NTG and TG-derived microbiomes were significantly different, and the estimated fraction of total variation explained by genotype (R2). At months 1 and 3 of age, there was no effect by genotype, neither qualitatively nor statistically (R2<2%, P>0.1). However, by 6 months of age, the microbial beta diversity of the TG and NTG had differed significantly from each other (R2=7%, P= 0.0001) and this genotype effect continued to diverge such that by month 12, the difference between TG vs. NTG-derived microbiomes accounted for 10% of the total variance (R2= 10%, P= 0.0001). Sensitivity analysis for litter-effect, using only 11 TG mice with their 22 NTG littermates, also showed no difference at 1 or 3 months, after which the genotypes begin to diverge progressively reaching significance at month 6 (R2= 5%, P= 0.02) and 12 (R2= 9%, P= 0.0009) **(Table 1b)**. We therefore conclude that the TG genotype, defined by α-syn overexpression, leads to an age-dependent shift in microbiome composition that differs from their NTG littermates.

**Table 1:** Testing for differences in the global composition of gut microbiome (beta-diversity) between Thy1-SNCA transgenic and non-transgenic mice Differences in the global composition of the gut microbiome between transgene (TG) and non-transgenic (NTG) mice were tested for using permutational analysis of variance (PERMANOVA). All tests were adjusted for total sequence count per sample. Tests at 1, 3 and 6 months were also adjusted for the first PC from PCA (see **Figure 1**). PERMANOVAs were performed once using all mice **(a)** and again using only mice from litters with both TG and NTG littermates **(b)**. R2, the proportion of inter-sample variation in microbiome composition explained by differences between TG vs NTG; F, pseudo-F statistic, the ratio between the amount of variance explained by differences between TG vs NTG and the residual variance in the data; P, permuted P-value from PERMANOVA computed using 9,999 permutations.

| Age (months) | a. All mice (N=54) |  |  | b. Litters with both TG and NTG (N=31) |  |  |
| --- | --- | --- | --- | --- | --- | --- |
|  | R2 | F | P | R2 | F | P |
| 1 | 0.02 | 1.49 | 0.13 | 0.03 | 1.53 | 0.15 |
| 3 | 0.02 | 1.53 | 0.12 | 0.03 | 1.69 | 0.1 |
| 6 | 0.07 | 5.03 | <1e-04 | 0.05 | 2.19 | 0.02 |
| 12 | 0.1 | 6.01 | <1e-04 | 0.09 | 2.97 | 5e-04 |

### α-syn overexpression results in reduced diversity in mouse gut microbiome

We next tested intra-individual variation (alpha-diversity) to determine whether the genotypes impacted the richness and evenness of the microbiome within an individual. We tested this alpha-diversity by three methods: species richness, Simpson, and Shannon indices **(Table 2)**. The Shannon index showed an age-dependent progressive reduction in diversity in TG as compared to NTG which reached statistical significance by month 12 (P = 0.005) and was confirmed in the sensitivity analysis of littermates. Decreases in alpha-diversity over time are often attributed to the loss of diversity, however it may also be interpreted as an inability to build diversity within the community. Nonetheless, as the α-syn overexpressing TG mice age, we observed significantly decreased diversity compared to their NTG littermates.

**Table 2:** Testing for differences in intra-sample gut microbiome diversity (alpha-diversity) between Thy1-SNCA transgenic and non-transgenic mice Alpha-diversity for each mouse was measured using Shannon, Simpson, and Richness diversity metrics. Differences in diversity metrics between transgene (TG) and non-transgenic (NTG) mice were tested for using linear regression. All tests were adjusted for total sequence count per sample. Tests at 1, 3 and 6 months were also adjusted for the first PC from PCA (see **Figure 1**). Tests were performed once using all mice **(a)** and again using only mice from litters with both TG and NTG littermates **(b)**. *t*, the standardized beta coefficient from linear regression (i.e., the difference in mean alpha-diversity metric between groups divided by its standard error); P, the raw P-value of the beta coefficient from linear regression.

| Age (months) | a. All mice (N=54) |  |  |  |  |  | b. Litters with both TG and NTG (N=31) |  |  |  |  |  |
| --- | --- | --- | --- | --- | --- | --- | --- | --- | --- | --- | --- | --- |
|  | Shannon |  | Simpson |  | Richness |  | Shannon |  | Simpson |  | Richness |  |
|  | t | P | t | P | t | P | t | P | t | P | t | P |
| 1 | -0.11 | 0.91 | 0.24 | 0.81 | -0.34 | 0.73 | 0.78 | 0.44 | 1.52 | 0.14 | -1.22 | 0.23 |
| 3 | -0.75 | 0.46 | -0.9 | 0.38 | 0.57 | 0.57 | -0.9 | 0.38 | -0.67 | 0.51 | -0.21 | 0.83 |
| 6 | -1.61 | 0.11 | -1.28 | 0.21 | 0 | 1 | -1.25 | 0.22 | -1 | 0.33 | -0.23 | 0.82 |
| 12 | -2.94 | 0.005 | -1.43 | 0.16 | -0.65 | 0.52 | -1.96 | 0.06 | -0.75 | 0.46 | -1.2 | 0.24 |

### Certain species of *Lactobacillus* and *Bifidobacterium* fail to bloom in α-syn overexpressing mice

Given our observations thus far, we have observed that TG and NTG mouse microbiomes diverge with age with the greatest difference occurring at 12 months of age. Our next step was to identify the specific bacterial species that drive the compositional shifts underlying this dysbiosis. We therefore designed a two-step analysis to specifically answer which taxa are driving the dysbiosis and when through the life of the mouse do these changes appear. First, we conducted a microbiome-wide association study (MWAS) where we tested the differential abundance of species and genera at month 12, when the divergence was the greatest, and identified taxa that were significantly altered between genotypes. Then, we longitudinally traced the relative abundances of the specified TG-associated taxa throughout aging at 1, 3, 6 and 12 months of age.

A total of 48 species and 29 genera were captured in the whole dataset after stringent MetaPhlan3 thresholds were applied. Of these, 32 species and 19 genera were represented within the 12 month aged dataset. We conducted species-level and genus-level MWAS, at this time point, once with all 54 mice (setting significance at FDR<0.05) and once with the 31 littermates (sensitivity analysis, requiring a trend as seen in 54 mice, and calling it significant at P<0.05). Species-level MWAS revealed 6 species that had significantly lower abundance in TG than in NTG (**Table 3a**). These included three species of *Lactobacillus*, each reduced by 14 to 50-fold in TG α-syn overexpressing mice as compared to NTG wildtype mice. Namely, *L. intestinalis* (fold change in TG vs. WT (FC)=0.13, FDR=1E-4), *L. reuteri* (FC=0.07, FDR=2E-3), and *L. johnsonii* (FC= 0.02, FDR= 2E-03) were substantially decreased. The three other species included, *Bifidobacterium pseudolongum* that was reduced by 20-fold (FC=0.05, FDR= 5E-4), *Muribaculum intestinale* with a 7-fold reduction (FC=0.05, FDR= 5E-4), and *Muribaculaceae bacterium DSM_10372* which was reduced by 5-fold (FC=0.20, FDR= 2E-3). All 6 of these species were further confirmed to be reduced in TG in sensitivity analysis of littermate (**Table 3b**). Therefore, we conclude that these taxa may be sensitive to the intestinal environment created by α-syn overexpression.

**Table 3:** Species and genera with significantly different abundances in Thy1-SNCA transgenic mice compared to non-transgenic mice Species- and genus-level differential abundance analysis between transgenic (TG) and non-transgenic (NTG) mice was performed on mouse samples collected at 12 months using linear regression with log2-transformed relative abundances as implemented in MaAsLin2. The analysis was adjusted for total sequence count per sample. Tests were performed once using all mice **(a)** and again using only mice from litters with both TG and NTG littermates **(b)**. Taxa whose relative abundance differed between TG and NTG groups at a multiple testing corrected significance of <0.05 when analyzing all mice are shown. Full results are provided in **Supplementary Tables 2** and **3**. N, number of mice in which taxa were detected; Mean RA, mean relative abundance; P, raw P-value from testing with MaAsLin2; FDR, false discovery rate q-value calculated via the Benjamini-Hochberg method; FC [95% CI], fold change and 95% confidence interval of the TG group relative abundance compared to the NTG group.

| Table 3. Species and genera with significantly different abundances in Thy1-SNCA transgenic mice compared to non-transgenic mice |  |  |  |  |  |  |  |  |  |  |  |  |  |
| --- | --- | --- | --- | --- | --- | --- | --- | --- | --- | --- | --- | --- | --- |
| Taxa | a. All mice (N=54) |  |  |  |  |  |  | b. Litters with both TG and NTG (N=31) |  |  |  |  |  |
|  | N TG | N NTG | Mean RA TG | Mean RA NTG | P | FDR | FC [95% CI] | N TG | N NTG | Mean RA TG | Mean RA NTG | P | FC [95% CI] |
| Species |  |  |  |  |  |  |  |  |  |  |  |  |  |
| *Bifidobacterium pseudolongum* | 3 | 21 | 0.1% | 14.8% | 3E-5 | 5E-4 | 0.05 [0.02; 0.19] | 1 | 13 | 0.1% | 10.5% | 5E-3 | 0.07 [0.01; 0.4] |
| *Lactobacillus intestinalis* | 5 | 26 | 1.1% | 5.0% | 5E-6 | 1E-4 | 0.13 [0.06; 0.29] | 2 | 17 | 1.1% | 5.2% | 1E-3 | 0.24 [0.11; 0.52] |
| *Lactobacillus johnsonii* | 13 | 29 | 2.5% | 9.3% | 4E-4 | 2E-3 | 0.08 [0.02; 0.29] | 7 | 18 | 3.3% | 8.7% | 0.03 | 0.11 [0.02; 0.71] |
| *Lactobacillus reuteri* | 13 | 30 | 6.2% | 12.2% | 3E-4 | 2E-3 | 0.07 [0.02; 0.27] | 7 | 19 | 7.7% | 12.3% | 0.01 | 0.08 [0.01; 0.49] |
| *Muribaculaceae bacterium DSM 103720* | 15 | 29 | 1.1% | 2.5% | 4E-4 | 2E-3 | 0.2 [0.09; 0.47] | 7 | 18 | 1.4% | 2.5% | 0.02 | 0.23 [0.07; 0.76] |
| *Muribaculum intestinale* | 16 | 31 | 1.8% | 2.7% | 3E-4 | 2E-3 | 0.14 [0.05; 0.38] | 8 | 19 | 2.8% | 2.8% | 0.02 | 0.15 [0.03; 0.68] |
| Genera |  |  |  |  |  |  |  |  |  |  |  |  |  |
| *Bifidobacterium* | 3 | 21 | 0.1% | 14.8% | 3E-5 | 6E-4 | 0.05 [0.02; 0.19] | 1 | 13 | 0.1% | 10.5% | 5E-3 | 0.07 [0.01; 0.4] |
| Unclassified *Muribaculaceae* | 15 | 29 | 1.1% | 2.5% | 4E-4 | 3E-3 | 0.2 [0.09; 0.47] | 7 | 18 | 1.4% | 2.5% | 0.02 | 0.23 [0.07; 0.76] |
| *Muribaculum* | 16 | 31 | 1.8% | 2.7% | 3E-4 | 3E-3 | 0.14 [0.05; 0.38] | 8 | 19 | 2.8% | 2.8% | 0.02 | 0.15 [0.03; 0.68] |

As suspected from species-level MWAS, genus-level MWAS identified *Bifidobacterium, Muribaculum*, and an unclassified genus in the *Muribaculaceae* family as being reduced in the α-syn TG mice (**Table 3**). Surprisingly, despite the reduction of three species of *Lactobacillus*, there was no signal for the overall *Lactobacillus* genus. Supplementary **Tables S2-S5** contain the full MWAS results. *Bifidobacterium, Muribaculum*, and the unclassified *Muribaculaceae* each had only one species from those genera detected in this dataset (**Table S2**), hence their relative abundances and fold changes were similar at both genus and species level. Conversely, the *Lactobacillus* genus contained four species: *L. intestinalis, L. reuteri, L. johnsonii*, and *L. murinus (***Table S2)**. The former three were individually significantly reduced in TG mice at 12 months of age. However, these species were relatively rare within the TG population, adding up to a combined relative abundance of only 9% in TG mice vs. 26% in NTG. In contrast, *L. murinus* is not only common, but it was also elevated in TG mice, albeit not significantly (mean relative abundance of 22% in WT and 42% in TG, FDR=0.15). Thus, when the four species are pooled together at the genus level, these differential abundances mask the effects at the species level (**Table S3)**. Hence, *Lactobacillus*, the only genus with multiple and the most significant signals at species level MWAS, was missed at genus level MWAS. Further, suggesting that distinct species of *Lactobacillus* may be selected for (*L. murinus*), or against (*L. intestinalis, L. reuteri, L. johnsonii*), in the intestinal environment created by α-syn overexpression over aging in these mice.

The differential abundances of these specific species could be due to their enrichment in the NTG mice over time, their loss in the α-syn overexpressing TG mice, or simply differential initial abundances that were maintained throughout age. To begin to differentiate between these possibilities, we tracked age-specific relative abundance profiles of the 6 species that characterized the dysbiosis at 12 months of age (**Figure 2, Table S6)**. Most striking was *Bifidobacterium pseudolongum*. In NTG mice, *B. pseudolongum* had a mean relative abundance of 2% at month 1, 6% at month 3, 8% at month 6, and 15% at month 12. In TG mice, however, *B. pseudolongum* constituted only 0.2% of microbiome at month 1, 0.8% in months 3 and 6, and 0.1% at month 12. Thus, TG were already depleted of *B. pseudolongum* by month 1, or it had never colonized effectively at birth. While NTG mice displayed a robust and progressive increase in the abundance of *B. pseudolongum* with age, this enrichment never occurred in the TG mice. Other TG-associated species do not show a significant difference at 1 or 3 months., maintaining their abundance at these early ages. The three *Lactobacillus* species diverge after month 3, and *Muribaculaceae and Muribaculum* species after month 6. Tracking the genera-level abundance by age would be identical to species-level tracking for all three TG-associated genera (*Bifidobacterium, Muribaculaceae and Muribaculum)* because only one species was detected in these three genera levels. Overall, these data demonstrate that α-syn overexpression creates an intestinal environment that limits the growth of certain species within the gut microbiome and prevents a more diverse community from forming with age.

**Figure 2.**
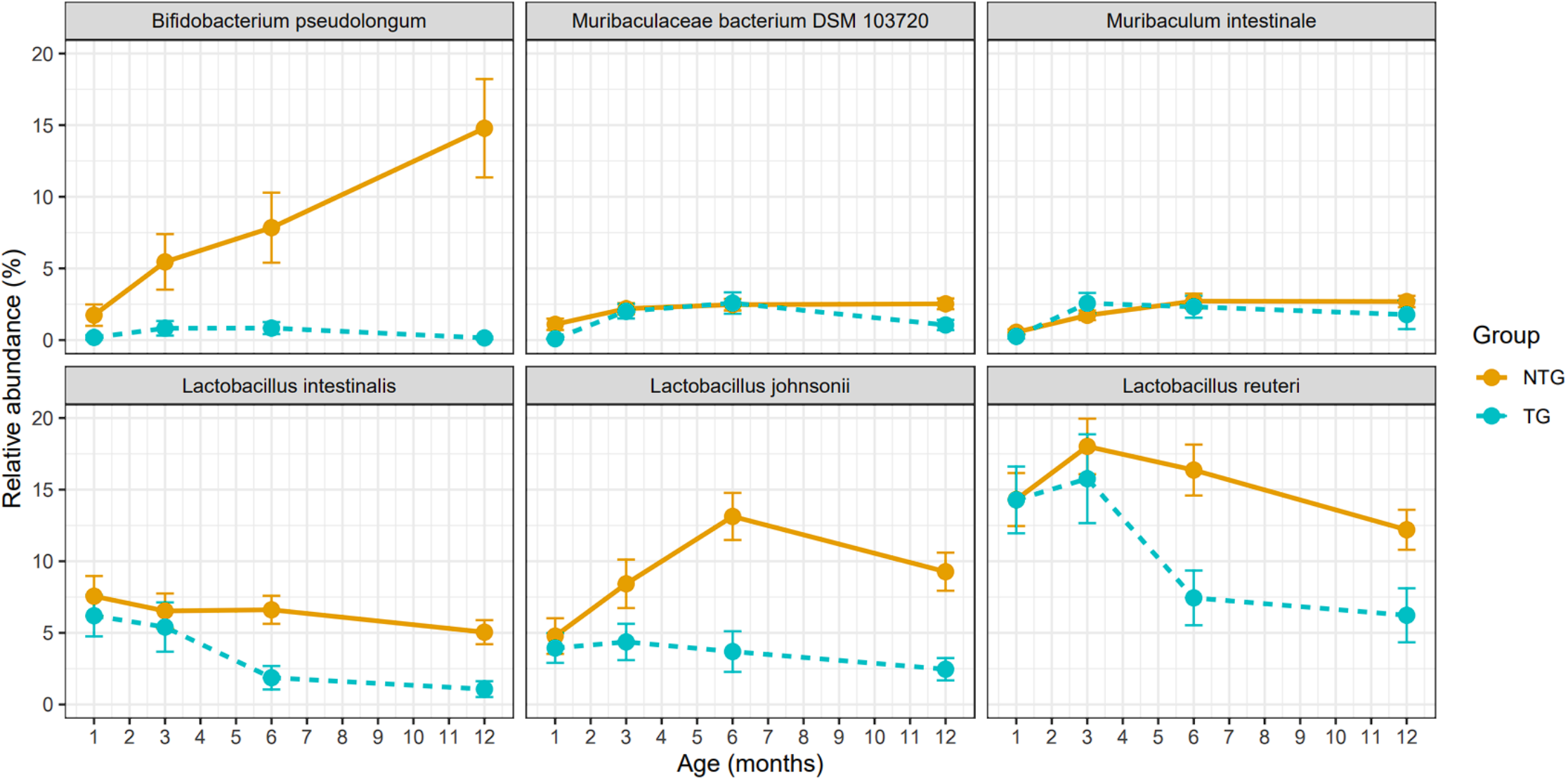
Changes over time in the relative abundance of species found differentially abundant between Thy1-SNCA transgenic and non-transgenic mice. Six species were found to be differentially abundant between 12-month-old Thy1-SNCA transgenic (TG, blue points and lines) and non-transgenic (NTG, orange points and lines) mice (Table 3), and their relative abundances were plotted over time to observe how they change with age. Each point represents the mean relative abundance of TG or NTG mice at a certain time point and error bars extending from each point represents the standard error of the mean. Differences in relative abundances at each time point between TG and NTG mice were formally tested using MaAsLin2 (Supplementary Table 6).

## DISCUSSION

In this longitudinal observational study, we assessed the impact of α-syn overexpression on the gut bacterial community in mice. We identified that α-syn overexpression drives an age-dependent dysbiosis in mice, characterized by diminished diversity and a decrease in specific *Lactobacillus* and *Bifidobacteria* species. With a growing consensus that distinct alterations to the gut microbiome are associated with PD, understanding how and when various models of PD-pathology recapitulate this aspect of disease is essential. The physiological impacts that might influence the GI environment during PD are complex and limit the ability to pinpoint which precise factors affect the microbiome composition and when in the disease process do these take place.

In the present study, we selected the Thy1-hSNCA (line 61) α-syn overexpressing mouse model as it has been fairly well-characterized in its pathological progression. In addition to α-syn pathological deposition in the CNS, mice display behavioral phenotypes, striatal dopamine depletion, and significant neuroinflammation in the absence of overt neurodegeneration (*21, 22, 27*). Line 61 mice express α-syn in the GI tract, and also display characteristic slowed GI motility as early as 2.5-3 months of age modeling the PD prodrome (*20*). This is notable, as we observe no genotype-dependent effect on the microbiome composition until after the 3 month time point. While not directly assessed, this suggests that α-syn overexpression induces GI deficits prior to the microbiome compositional shift in this model, as others observe GI dysfunctions arising concurrent and prior to this age (*20, 28*). It may be that a slowed intestinal environment selects against those taxa we observe increased with age in the NTG controls. Understanding how slowed GI motility directly impacts microbiome composition in this and other models will provide insight into shared microbiome features across neurodegenerative diseases.

This α-syn overexpression model also begins to show mild to moderate increases in inflammatory markers beginning at 2 months of age and are subsequently exacerbated by 6 months of age (*27*). Early changes include microgliosis, astrogliosis, and increased expression of the pro-inflammatory cytokine tumor necrosis factor (TNF) and the pattern recognition receptor Toll-like receptor 2 (TLR) (*27*). Given a potential role for α-syn in immune activation, particularly via the interaction with TLR2 to mediate myeloid activation (*29*), altered immune signaling in the GI tract may contribute alongside slowed motility to induce a microbiome compositional shift. This compositional shift may then feed back onto the immune system thus exacerbating inflammation in a pathological feed-forward cycle. Importantly, the compositional shift in the microbiome appears well before dopamine signaling is diminished in this model (∼14 months of age) (*22*). Thus, the microbiome compositional shifts we observe may be a harbinger of pre-manifest PD deficits and reflect a secondary pathology of α-syn overexpression.

In this dataset, α-syn overexpressing mice harbor decreased levels of *Lactobacillus* and *Bifidobacteria* species and do not obtain a diverse microbiome in adulthood. Two prior studies have conducted low resolution 16S profiling to explore the effects of transgenic α-syn production on the microbiome (*15, 19*). Despite differences in our sample sizes (5-10 vs. 54), genotypes (wild-type vs. α-syn), and sequencing method (16S vs. high resolution shotgun metagenomics), we note a few key shared features. All studies clearly demonstrate that α-syn overexpression leads to microbiome compositional changes that manifest shortly after birth and accelerate with age. The precise species that are selected for or against within each study may be secondary, driven by mouse strain, specific α-syn transgene, the microbiome facility, and other extrinsic factors. Each study also demonstrates that α-syn transgenic mice have a less diverse microbiome than wild-type controls. While Radisavljevic et al. attribute this to a loss of diversity, our data suggest instead it is an inability to gain diversity from a simpler newborn microbiome(*30*). It is noteworthy that *Lactobacillus* and *Bifidobacteria* emerge as the most significantly aberrant in both human PD and α-syn rodent models, suggesting they are sensitive to a feature with the PD-associated intestinal environment. While unexpected differences in the trends of these specific species appear, it underscores the importance of these two taxa with the intestinal environment shaped by α-syn and Parkinson’s disease. Given emerging data highlighting contributions of various bacterial species to promoting inflammation and synuclein pathology (*18, 20, 31*), understanding why certain microorganisms come to be enriched in the PD-associated microbiome is essential.

The more intangible challenge is extrapolating phenotypes in this mouse model and drawing parallels with human PD. Setting aside the obvious biological differences between mice and humans, no existing rodent model fully recapitulates all attributes of PD. However, the specific aspects of how each model system manifests, and if/how the microbiome composition changes within them, would allow an understanding of what drives the microbiome compositional shifts in PD. Importantly, in this study, we are assessing a model which overproduces α-syn pan-neuronally prior to birth. It is not known if people who develop sporadic PD congenitally overexpress *SNCA*, and if not, when in life α-syn accumulation is triggered. Nonetheless, this mouse model provides a powerful tool for investigating the underlying mechanisms of disease. In particular investigation of synuclein pathologies in pre-manifest PD and how they may align with compositional shifts in the gut microbiome. Identification of those early compositional shifts as they align with key stages of pre-manifest PD and its progression could inform microbiome-based diagnostics and prognostic indicators in humans. More coercively, these earliest microbiome compositional differences may underlie differences in future disease outcomes. If human studies are any indication, our present results are only the tip of the iceberg in delineating the elusive question of when and how the PD-associated microbiome arises.

## Conclusion

Overall, in mice overexpressing α-syn, we find significant, age and genotype dependent bacterial taxa that become altered over age. Futhermore, we found that α-syn overexpression can drive alterations to the gut microbiome composition and suggest that it limits the expansion of diversity with age. Given emerging data on potential contributions of the gut microbiome to PD pathologies, our data provide an experimental foundation to understand how the PD-associated microbiome may arise as a trigger or co-pathology to disease.

**Supplementary Figure 1.**
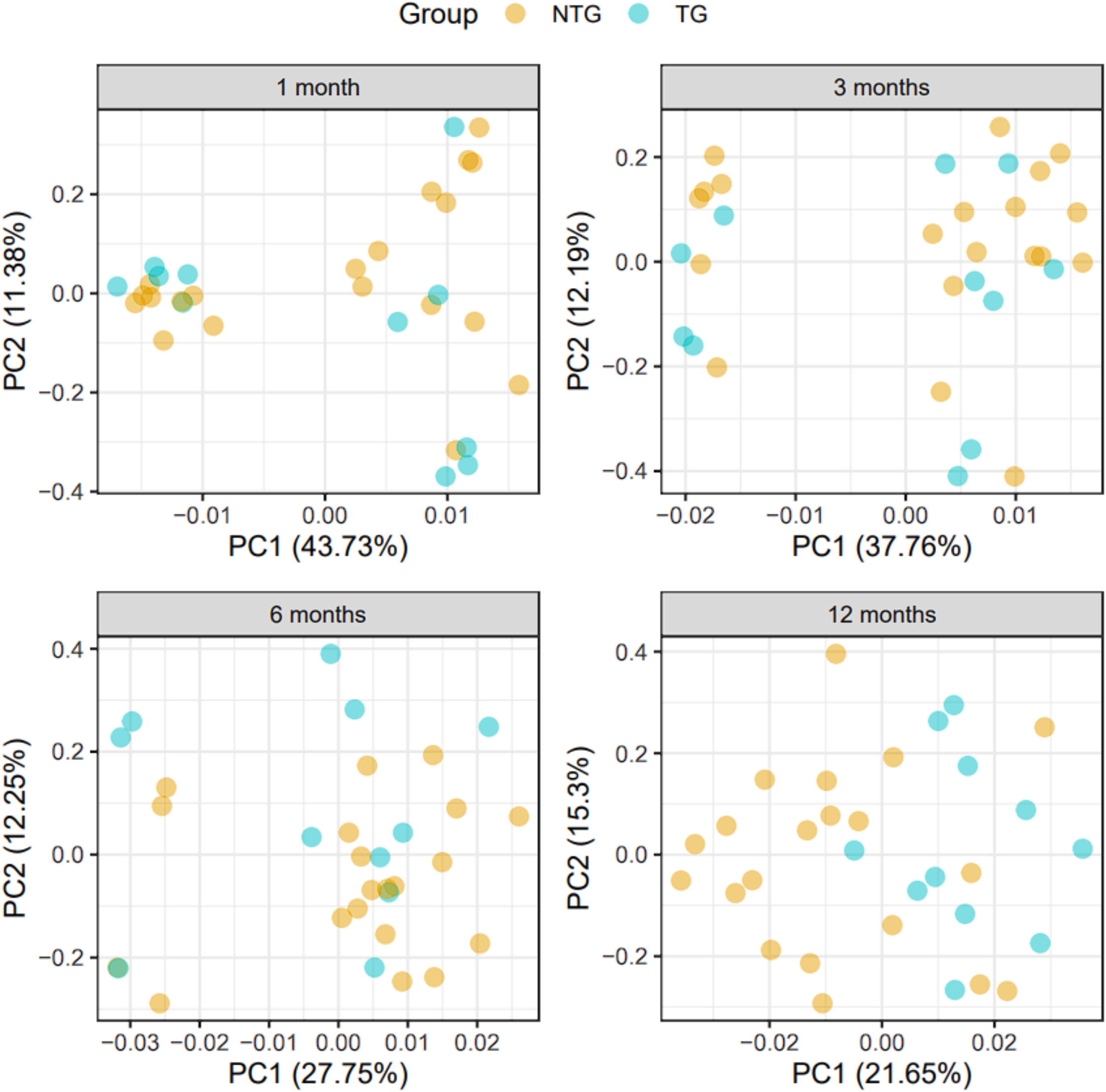
Inter-sample differences in gut microbiome composition (beta-diversity) of Thy1-SNCA transgenic and non-transgenic mice at 1, 3, 6 and 12 months including only mice from litters with both transgenic and non-transgenic litter mates. To ensure principal component (PC) analysis results were robust to litter effects, PC analysis was performed again to visualize inter-sample differences in gut microbiome composition (beta-diversity) between Thy1-SNCA transgenic (TG, blue points, N = 11) and non-transgenic (NTG, orange points, N = 20) mice, including only mice who came from litters with both TG and NTG littermates. Each point in the plots represents the composition of the gut microbiome of one unique mouse sample at a certain time point and distances between points indicate degree of similarity of a mouse sample to others. Percentages on the x- and y-axis correspond to the percent variation in gut microbiome compositions explained by PC1 and PC2. The differences between TG and NTG groups for each time point were formally tested using PERMANOVA (Table 1b).

**Supplementary Figure 2.**
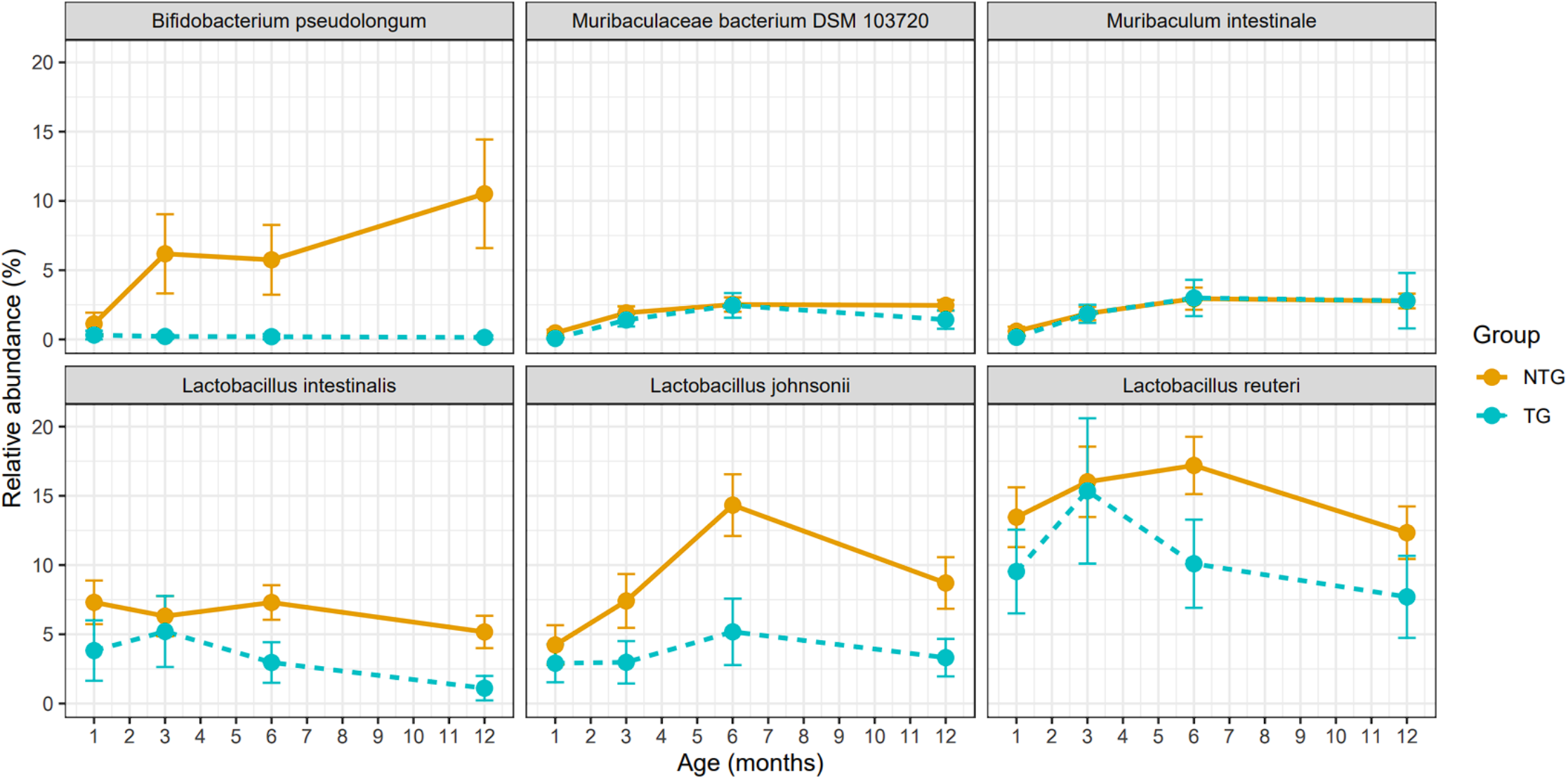
Changes over time in the relative abundance of species found differentially abundant between Thy1-SNCA transgenic and non-transgenic mice including only mice from litters with both transgenic and non-transgenic litter mates. Changes in relative abundance over time for the six differentially abundant species were visualized again including only mice from litters with both TG and NTG littermates. Each point represents the mean relative abundance of TG or NTG mice at a certain time point and error bars extending from each point represents the standard error of the mean.

## Acknowledgements

This work was supported by Parkinson Association of Alabama Research Acceleration Fund (to HP), Aligning Science Across Parkinson’s [ASAP-020527 to TRS and HP, ASAP-000375 to AH] through the Michael J. Fox Foundation for Parkinson’s Research (MJFF). For the purpose of open access, the authors have applied a CC BY public copyright license to all Author Accepted Manuscripts arising from this submission. Opinions, interpretations, conclusions, and recommendations are those of the authors and are not necessarily endorsed by the funding agencies.

